# RNAStructViz: Graphical base pairing analysis

**DOI:** 10.1101/2021.01.20.427505

**Authors:** Maxie Dion Schmidt, Anna Kirkpatrick, Christine Heitsch

## Abstract

**Summary:** We present a new graphical tool for RNA secondary structure analysis. The central feature is the ability to visually compare/contrast up to three base pairing configurations for a given sequence in a compact, standardized circular arc diagram layout. This is complemented by a built-in CT-style file viewer and radial layout substructure viewer which are directly linked to the arc diagram window via the zoom selection tool. Additional functionality includes the computation of some numerical information, and the ability to export images and data for later use. This tool should be of use to researchers seeking to better understand similarities and differences between structural alternatives for an RNA sequence.

**Availability and implementation:** https://github.com/gtDMMB/RNAStructViz/wiki

**Author contacts:**, and

## 1 Introduction

As the cellular roles for RNA molecules continue to grow [2], so does the importance of gaining functional insight from structural analyses. While obtaining the 1D sequence information is now relatively easy, characterizing 3D molecular conformations is still comparatively hard. Hence, understanding the 2D secondary structures, i.e. the intra-sequence base pairing, remains a crucial component of ribonomics research.

There are many programs which generate possible secondary structures for a given RNA sequence. However, understanding the similarities and differences between predictions can be a challenge. We present here a new graphical tool, RNAStructViz, which can visualize up to three different configurations simultaneously. In moving beyond pairwise comparisons, there is greater potential for insight from spotting structural patterns.

## 2 Core features

As illustrated in Figure 1, RNAStructViz has a main navigator window where the uploaded secondary structures are automatically organized into folders by common sequence. Once a folder has been selected from the left half of the main window, the “View Diagram” and “Statistics” windows for that sequence can be opened from the right half. Multiple windows for the same sequence or for different sequences can be open concurrently.

**Fig. 1:**
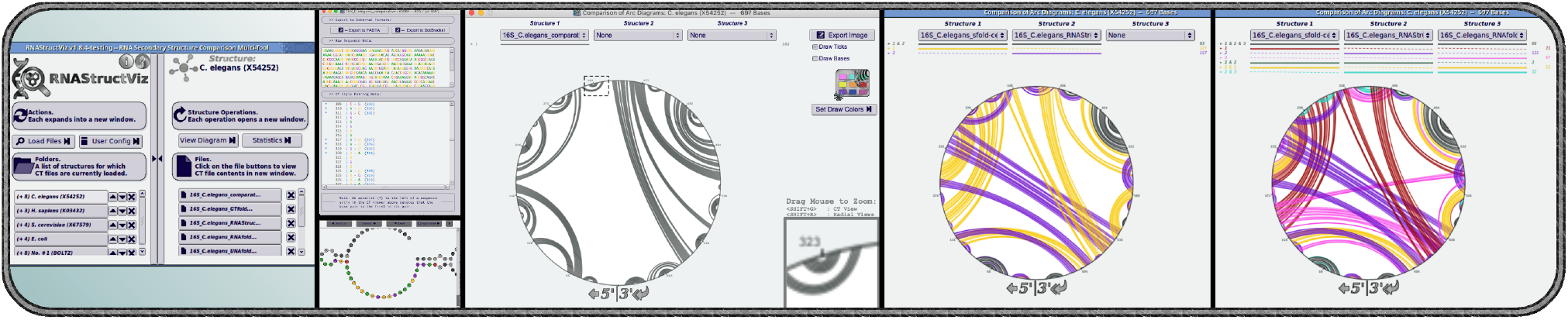
Visual analysis of RNA secondary structures. Five panels show screenshots for a 16S ribosomal RNA sequence with 697 nucleotides. The main navigation window (1st) is followed by a file viewer listing sequence and base pairs (2nd top) and radial substructure display (2nd bottom). These features linked to the zoom tool (3rd, dashed box & bottom right) in the arc diagram display. Up to 3 secondary structures for the same sequence can be viewed simultaneously. When 2 or 3 are selected (4th, resp. 5th), the common base pairs are distinguished by colors, identified in the legend at the top along with the counts. In these ways, RNAStructViz facilitates visual analysis of similarities and differences among structural alternatives.

Within a diagram window, users have the option of selecting up to three secondary structures to view simultaneously. The structures are drawn as circular arc diagrams; this compact, standardized layout permits visual inspection of similarities and differences even for longer sequences. Each subset of common base pairs is highlighted in a different color, identified in the window legend. Like a Venn diagram, useful comparisons are limited to *n* = 2 or 3 structures since the number of subsets/colors grows as 2^*n*^.

Of particular note (c.f. Fig 1, central panel, bottom right corner) is the zoom tool which permits detailed inspection of selected base pairs in two ways. First, a CT-style file viewer can be opened at that point in the sequence. (The file viewer is also accessible from within a sequence folder on the right of the main window.) Second, a radial layout of the pairings, and adjacent substructure, can be viewed. Both options greatly increase the level of resolution of structural information available for visual analysis. We also note that both the arc and radial diagrams can be exported as PNG images or in SVG format for later use.

The statistics window provides numerical information, such as the types of base pairs in each structure. It also calculates one-to-many pairwise comparisons between sets of base pairs; for a selected reference *A*, the number of “true positive” (*A* ∩ *B*), “false positive” (*B* \ *A*) and “false negative” (*A* \ *B*) pairings are tabulated for all other structures B. If desired, this data is used to compute sensitivity and selectivity (a modified PPV [4]), which are displayed in a ROC plot. Finally, all numerical information is available in a single table that is exportable as a CSV file.

## 3 Comparison with related software

Other freely available tools for RNA secondary structure graphical analysis include FORNA [5], jViz.RNA [8]. R-chie [6], RNAbows [1], and VARNA [3]. Table 1 lists some key features in four categories, and indicates their support in RNAStructViz and these related programs.

**Table 1.** A comparison of selected features across related tools; an extended survey appears in the wiki. (*) Although FORNA can display multiple structures in the same window, the similarities/differences are not highlighted. (**) Some versions of jViz.RNA provide statistical comparisons.

|  | RNAstructViz | FORNA | jViz.RNA | R-chie | RNAbows | VARNA |
| --- | --- | --- | --- | --- | --- | --- |
| ↓ Feature sets | Software support → |  |  |  |  |  |
| ① — Platform and availability — |  |  |  |  |  |  |
| Mac OSX support | ✓ | ✓ | ✓ | ✓ | ✓ | ✓ |
| Linux / Unix support | ✓ | ✓ | ✓ | ✓ | ✓ | ✓ |
| Windows support | ✗ | ✓ | ✓ | ✓ | ✓ | ✓ |
| Open source software | ✓ | ✓ | ✓ | ✓ | ✗ | ✓ |
| Requires external libraries | ✓ | ✓ | ✓ | ✓ | ✗ | ✓ |
| ② — Software usability criteria — |  |  |  |  |  |  |
| Graphical user interface | ✓ | ✓ | ✓ | ✓ | ✓ | ✓ |
| Web interface | ✗ | ✓ | ✗ | ✓ | ✓ | ✗ |
| Multi-window interface | ✓ | ✗ | ✗ | ✗ | ✗ | ✗ |
| Compares 2 structures at once | ✓ | ✓ | ✗ | ✓ | ✓ | ✗ |
| Compares 3 structures at once | ✓ | ✓* | ✗ | ✗ | ✗ | ✗ |
| ③ — Support for standard formats — |  |  |  |  |  |  |
| CT files | ✓ | ✗ | ✓ | ✓ | ✗ | ✓ |
| Dot-bracket files | ✓ | ✗ | ✓ | ✓ | ✗ | ✓ |
| Built-in file viewer | ✓ | ✗ | ✗ | ✗ | ✗ | ✓ |
| Requires specialized format | ✗ | ✓ | ✗ | ✗ | ✗ | ✗ |
| Can edit sequence data | ✗ | ✓ | ✓ | ✗ | ✓ | ✓ |
| ④ — Views and diagram type support — |  |  |  |  |  |  |
| Has comparison statistics | ✓ | ✗ | ✓** | ✓ | ✓ | ✗ |
| Plots circular arc diagrams | ✓ | ✗ | ✓ | ✓ | ✓ | ✗ |
| Plots radial diagrams | ✓ | ✓ | ✓ | ✗ | ✗ | ✓ |

RNAStructViz is a useful addition to this set of tools since it supports both a multi-window interface and automatic triple structural comparison. Both features should enable researchers to better understand the similarities and differences between alternative RNA secondary structures.

## 4 Availability and installation

RNAStructViz is open source, and available on Github.com under a GPL-V3 license. The project repo also includes a detailed wiki. The program has been tested on Mac OSX (10.14 – Mojave) and Linux (CentOS and Ubuntu 16.04 32-bit, 18.04 64-bit). For Mac users, a Homebrew package is provided to simplify installation. If installing from source, four C++ libraries are needed: OpenSSL for string hashing, Boost for cross-platform filesystem operations, FLTK for the lightweight GUI framework, and Cairo for enhanced graphics support. Additionally, the ViennaRNA toolkit [7] is needed to generate radial layouts of selected substructures.

## 6 Acknowledgements

Many collaborators have contributed to this project over the years; a full list of credits is given in the wiki. We particularly appreciated receiving testing feedback from (in alphabetical order): the Aviran Lab at UC Davis, the Laederach Lab at UNC Chapel Hill, and the Mathews Lab at U Rochester.

